## Supplementary material for "Atypical Structural Tendencies Among Low-Complexity Domains in the Protein Data Bank Proteome": File S3

**Supplementary Discussion**

The description here pertains to Figs S9-S10, which require more lengthy discussion to fully appreciate.

The analyses in Fig 7 and Figs S4-S6 suggest a general decline in the performance of secondary structure propensity scales at compositional extremes among LCDs. However, this does not indicate the degree of single-amino acid enrichment at which this occurs for each type of LCD, nor does it indicate how the residuals vary as a function of composition. Therefore, in order to thoroughly examine the relationships between the degree of enrichment for each amino acid within the various LCD types and the secondary structure propensity values from established scales, we decomposed LCDs into their composite “residue count” bins and performed regression analyses separately for each bin. For example, for each amino acid, all LCDs containing exactly 3 of that particular amino acid (out of 12) were parsed into a single bin/group. Once these groups were generated for each amino acid, the fraction of residues in α-helices or β-sheets was calculated within each group. This process was repeated independently for each of the “residue count” bins, from 3 residues corresponding to a single amino acid up through 12 residues corresponding to a single amino acid.

For each residue count bin, the average residual value from the highly-enriched LCD regression lines (from Figs S4-S5) was calculated across all α-helix or β-sheet propensity scales separately for each combination of residue count bin and LCD class (i.e. LCDs parsed by predominant amino acid). For example, for all LCDs containing exactly 3 alanine residues, the difference between the observed fraction of residues in α-helices among LCDs with 3 alanine residues *vis-à-vis* the highly-enriched LCD regression line was calculated for each of the propensity scales, then averaged across scales and stored as a single value. This was repeated for all combinations of LCD type and residue count bin, for both α-helices and β-sheets. This statistic effectively represents a comparison of LCDs within each “residue count” bin with the corresponding combined set of highly-enriched LCDs (i.e. LCDs that contain a single type of amino acid at ≥50% composition). The overall trend in these average residual values indicates the relationship between the observed secondary structure content and how far away it is from expectations based on propensity scales *as a function of the degree of enrichment for each amino acid (i.e. composition)*.

Fig S9 and Fig S10 indicate the average residual values for α-helix and β-sheet propensity scales, respectively. For reference, a residue count bin with a mean residual of 0.0 indicates that the average fraction of secondary structure corresponding to all protein segments in that residue count bin does not differ from the average fraction of secondary structure predicted by the regression line. Generally, each type of LCD falls into one of two categories: 1) those with mean residual that becomes more negative as single amino acid composition increases (indicating that the observed fraction of secondary structure becomes progressively *lower* than expected as a function of composition), and 2) those with mean residual that becomes more positive as single amino acid composition increases (indicating that the observed fraction of secondary structure becomes progressively *higher* than expected as a function of composition).

For α-helices, A-, D-, E-, G-, H-, K-, L-, N-, Q-, R-, S-, T-, W-, and Y-rich LCDs exhibit a decline in mean residual as composition increases, albeit with different curve shapes and slopes, which indicates the effect that each additional residue has on the relationship between observed and expected secondary structure content. It is worth noting that the sample size in each residue count bin generally decreases as residue count increases (refer back to Fig 1), so the final point in each case may be substantially affected by small sample size (e.g. as observed for L-, I-, and M-rich LCDs). For ease of interpretation, the size and opacity of each point (as well as the opacity of the preceding line segment) correspond to the LCD sample size in each bin. By contrast, F-, and I-rich LCDs generally maintain a positive mean residual that increases as residue count increases, with the exception of the last point, which corresponds to very low sample sizes in both cases. Additionally V-rich LCDs exhibit a general increase in mean residual when residue count is >6. Therefore, these LCDs appear to become increasingly more α-helical than expected as residue count increases, compared with established α-helix propensity scales.

For β-sheets, the mean residual for C-, D-, E-, F-, G-, K-, L-, N-, Q-, R-, S-, and T-rich LCDs tends to decrease as residue count increases, while the mean residual for V-, W-, and Y-rich LCDs tends to increase as residue count increases. Interestingly, for LCDs exhibiting an overall trend toward more negative mean residuals, most exhibit a slight increase in mean residual at residue counts around 4-6 before a precipitous decline, indicating that the decline in residual values only becomes apparent at very high levels of single amino acid enrichment.

Overall, these observations indicate a general decline in the efficacy of secondary structure propensity scales for LCDs, and highlight which residues most strongly contribute to this decline, as well as their composition dependencies.
